# Altered dorsal CA1 neuronal population coding in the APP/PS1 mouse model of Alzheimer’s disease

**DOI:** 10.1101/674374

**Authors:** Udaysankar Chockanathan, Emily Warner, Loel Turpin, M. Kerry O’Banion, Krishnan Padmanabhan

## Abstract

While the link between amyloid β (Aβ) accumulation and synaptic degradation in Alzheimer’s disease (AD) is known, the consequences of this pathology on coding remain unknown. We found that the entropy across neural ensembles was lower in the CA1 region in the APP/PS1 mouse model of Aβ, thereby reducing the population’s coding capacity. Our results reveal a network level signature of the deficits Aβ accumulation causes to the computations performed by neural circuits.

## Main Text

Alzheimer’s disease (AD) is a progressive neurodegenerative disorder associated with cognitive decline that is thought to arise in part due to the pathological accumulation of amyloid β (Aβ) plaques^1^ throughout the neocortex and hippocampus. Plaques cause a constellation of changes in neural circuits including, but not limited to, degradation of dendritic spines^2^, reductions in synapse density^2,3^, and increases in the intrinsic excitability of neurons^3^. Aβ pathology has been linked to various behavioral and cognitive changes^4,5^; for example, plaque burden correlates with degradation of place fields in the dorsal CA1 (dCA1) subfield of the hippocampus resulting in poor performance on spatial memory tasks^4^. Such behaviors require the orchestration of activity across large groups of neurons, or ensembles, whose dynamics are governed by the structure of neural circuits^6^. However, although Aβ pathology disrupts multiple features of these circuits^2,7,8^, the net effect of these changes on the structure of population activity and the resulting disruptions in neural computation remain unknown.

To address this question, we performed electrophysiological recordings in the hippocampus of awake APP/PS1 mice (model of Aβ pathology^9^, **Fig. 1a, d**), where dense amyloid plaques can be seen at 12 months of age^10^ and correspond to poor performance on spatial cognition and memory tasks, such as the T-maze alternation task^4,5^. High-density 128 channel arrays were targeted to the dCA1 region in head-fixed APP/PS1 and non-transgenic control mice (N = 4 male APP/PS1, N = 4 male control, age = 12-13 months, see **Supplemental Methods** for details) trained to run freely on a one-dimensional wheel. Up to 134 well-isolated single units were identified in each animal^11^ (control = 58±37 units, APP/PS1 = 59±50 units, see supplemental methods, **Fig. 1b-c, e-f, S1**). Representative rasters from a control (**Fig. 1g**) and APP/PS1 mouse (**Fig. 1h**) illustrate the complexity of patterns of ensemble activity across both groups.

**Figure 1.**
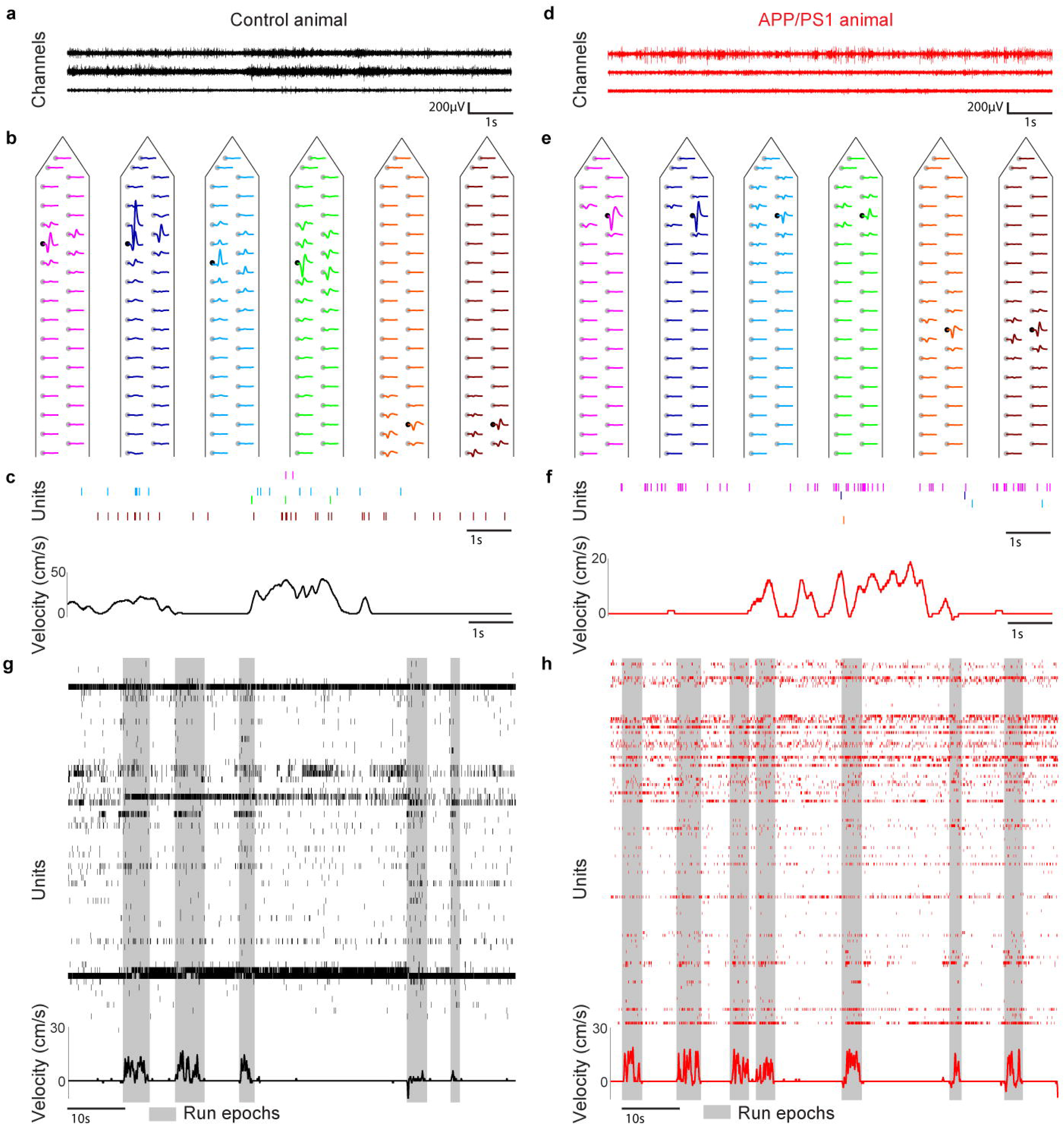
High-density awake recordings of dCA1 neuronal populations. (**a.d**) 10-second interval of raw electrophysiology data from three channels. (**b.e**) Mean waveform of each unit from channels in **a** and **d** as it appeared on the channel on which it was detected (black circle) and on the other channels on the electrode shank (grey circles). (**c,f**) Raster plot of sorted units and running velocity. Each tick denotes an action potential. (**g,h**) Raster plot of all units isolated in a representative control (**g**) and APP/PS1 (**h**) animal and simultaneous running behavior.

To characterize the statistics of this activity across ensembles in both control and APP/PS1 mice, we first calculated the entropy, a measure of the diversity of patterns of firing that occurred over the duration of the recording^12,13^ (**Fig. 2a, S2**).

**Figure 2.**
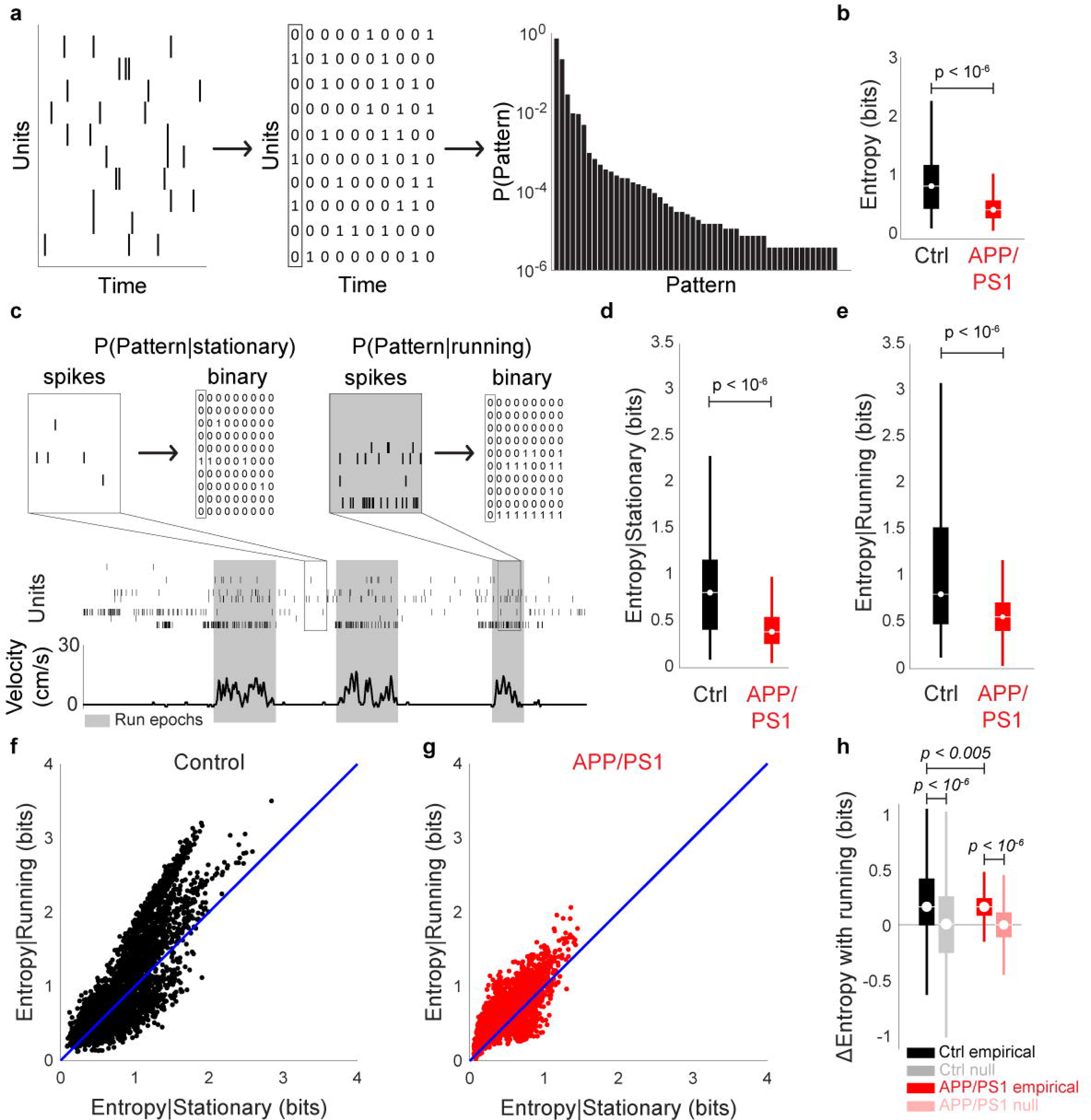
Entropy of dCA1 neuronal populations **(a)** Schematic of entropy calculation. **(b)** The APP/PS1 group had significantly lower entropy than controls. **(c)** Schematic of entropy calculation conditioned on behavioral state. **(d,e)** For both stationary (**d**) and running **(e)** conditions, the entropy of APP/PS1 animals was lower than that of controls. **(f,g)** Comparison of entropy for stationary and running entropy. **(h)** Entropy change with running. Null distributions were generated from a Gaussian centered at 0 with the same standard deviation as the corresponding empirical distributions. Both control and APP/PS1 animals exhibited a significant increase in entropy with running. However, the magnitude of this increase was larger in controls than APP/PS1 animals.

First, in dCA1, we observed that entropy in APP/PS1 animals was decreased relative to that of controls (control = 0.84±0.45 bits, APP/PS1 = 0.45±0.24 bits, *p* < 10^-6^, one-sided Wilcoxon rank-sum test, **Fig. 2b**), suggesting that the number of different patterns in APP/PS1 animals was smaller than in controls. The reduced entropy in APP/PS1 animals was significant across a range of ensemble sizes varying from 3 to 19 neurons (**Fig. S3, S4**), and was even preserved after normalizing entropy by firing rate (**Fig. S5**).

Different behavioral states, such as running or remaining stationary, impact features of both single neuron and population activity^14–16^. To understand how these behavioral states affected entropy in control and APP/PS1 mice, the frequency of different patterns of activity was estimated when animals were running and compared to when animals remained stationary (**Fig. 2c**). First, we found that running increased entropy in both control (stationary = 0.83±0.45 bits, running = 1.04±0.68 bits, *p* < 10^-6^, one-sided Wilcoxon rank-sum test, **Fig. 2f**,**h**) and APP/PS1 mice (stationary = 0.43±0.24 bits, running = 0.59±0.26 bits, *p* < 10^-6^, one-sided Wilcoxon rank-sum test, **Fig. 2g,h**). Interestingly, independent of whether the animal was stationary (control = 0.83±0.45 bits, APP/PS1 = 0.43±0.24 bits, *p* < 10^-6^, one-sided Wilcoxon rank-sum test) or running (control = 1.04±0.68 bits, APP/PS1 = 0.59±0.26 bits, *p* < 10^-6^, one-sided Wilcoxon rank-sum test), entropy was always lower in APP/PS1 animals than controls (**Fig. 2d,e, S6**-**7**), suggesting that the reduction in the number of possible network patterns was a general feature of ensemble activity. Furthermore, the magnitude of the change in entropy from stationary to running was significantly smaller in APP/PS1 mice than controls. (control = 0.21±0.38 bits, APP/PS1 = 0.16±0.17 bits, *p* < 0.005, one-sided Wilcoxon rank-sum test, **Fig. 2h**). Aβ pathology thus reduced not only the number of patterns generated by neural ensembles, but also the flexibility of those patterns across behaviors.

While the changes in entropy observed in APP/PS1 animals suggested that the diversity of network patterns was reduced with amyloid deposition, it remained unclear why such a reduction occurred. For instance, hypersynchronous neuronal activity, associated with the increased risk of seizures in human AD as well as mouse models^17,18^, could reduce entropy by increasing the occurrence of patterns of highly correlated neurons. By contrast, a similar, albeit mechanistically distinct reduction in entropy could occur due to the synapse loss and compromised dendritic structure seen in APP/PS1 animals^3^, which would result in fewer network patterns.

To disambiguate these different possibilities, we linked the statistics of network patterns to the functional coupling of neurons using maximum-entropy models that aim to predict patterns of activity with as few *a priori* assumptions of structure as possible^13,16,19^. For each ensemble, we fit both an independent firing model that only contained a term for the activity of each neuron (**h**_i_) and a pairwise interaction model that contained the **h**_i_ term for each neuron as well as a term for the functional coupling between pairs of neurons (**J**_ij_) (**Fig. S8**). This allowed us to estimate how cell autonomous properties, such as intrinsic excitability, and cell non-autonomous properties, such as pairwise interactions, shaped the patterns of the network, and, in turn, the entropy. To visualize this, we plotted the predicted patterns from the models and the empirical patterns for a representative control (**Fig. 3a,b**) and APP/PS1 (**Fig. 3c,d**) animal. Each point represents a different pattern and the color denotes the number of active units in that pattern. To quantify the goodness of fits between model and data, we used a measure of the distance between two probability distributions, the Kullbeck-Liebler Divergence (KLD). First, the KLD of the independent firing model was significantly larger for controls than APP/PS1 animals (control = 1.97×10^-2^±3.39×10^-2^, APP/PS1 = 3.81×10^-3^±6.29×10^-3^, *p* < 10^-6^, one-sided Wilcoxon rank-sum test, **Fig. 3e, S9, S10**), showing that a first order maximum-entropy model better predicted patterns of neuronal activity for APP/PS1 animals than for controls. Additionally, we found that the pairwise interactions model was also better at predicting patterns in APP/PS1 animals than controls (control = 9.17×10^-4^±1.20×10^-3^, APP/PS1 = 2.24×10^-4^±2.88×10^-4^, *p* < 10^-6^, one-sided Wilcoxon rank-sum test, **Fig. 3e, S11**). Interestingly, the **J**_ij_ terms in APP/PS1 animals were decreased compared to control animals (control = 0.02±1.10, APP/PS1 = −0.28±1.01, *p* < 0.05, one-sided Wilcoxon rank-sum test, **Fig. 3g, S12**), suggesting that a model incorporating reductions in functional coupling between neurons, possibly arising from reductions in dendritic length and branching^3^ or decreased synaptic density^2^, was better able to predict the paucity of dCA1 network patterns (**Fig. S13, S14**).

**Figure 3.**
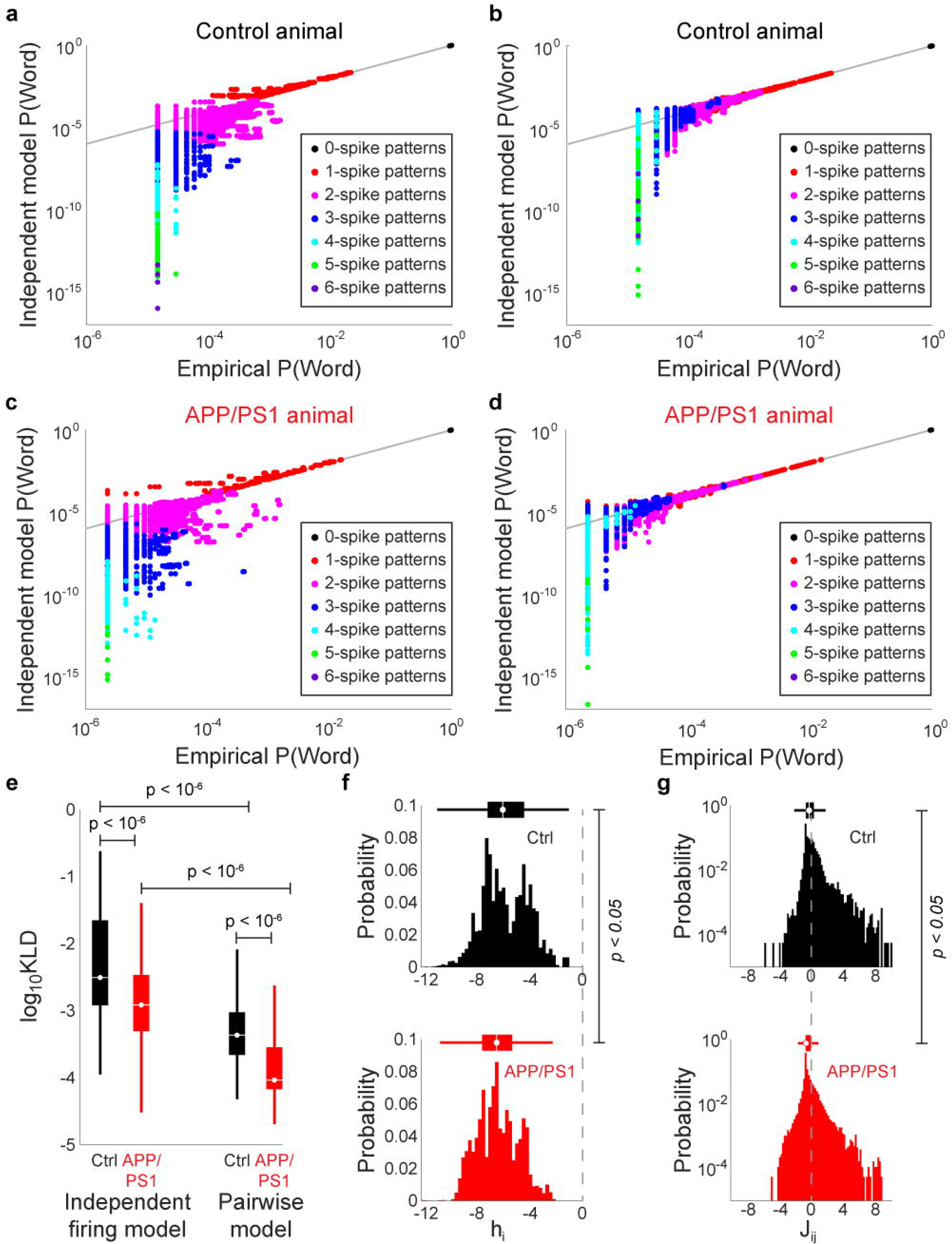
Maximum entropy models of dCA1 neuronal populations in control and APP/PS1 mice (**a-d**) Comparison of predicted and model word probabilities for a representative control (**a,b**) and APP/PS1 (**c,d**) animal for the independent model (**a,c**) and pairwise model (**b,d**) Colors denote the number of coactive neurons in each word. (**e**) Kullback-Liebler divergence (KLD) between empirical and model word probability distributions. For both models, the KLD nwas smaller for APP/PS1 mice than controls. Moreover, the pairwise models for both control and APP/PS1 groups had a lower KLD than the corresponding independent models. (**f,g**) Histograms of (**f**) h_i_ and (**g**) J_ij_ terms. Dashed line denotes 0. Both h_i_ and J_ij_ terms were smaller in the APP/PS1 group than the controls.

In summary, the decreased entropy observed in APP/PS1 animals revealed a reduction in the diversity of network patterns available to populations of neurons in dCA1. Such a decrease effectively constrains the number of patterns available to represent sensory stimuli or experiences, and this reduction in coding vocabulary could lead to the cognitive and spatial memory impairments seen in APP/PS1 mice^5^ and may provide clues into impairments in human AD^20,21^.

Additionally, we found that maximum-entropy models, both the independent and pairwise models, were better at predicting dCA1 population activity in APP/PS1 animals than controls. While these models do not explicitly reflect circuit level deficits such as synaptic connectivity, they provide insight into how the constellation of cellular and molecular changes in APP/PS1 and related models of AD may result in diminished coding capacity and network function^2,3^. The counterintuitive result that the pairwise interactions model was better able to account for dCA1 activity in APP/PS1 animals than controls suggests that once a change in functional coupling was accounted for, there was sufficient information to describe the diversity of network patterns in APP/PS1 animals. By contrast, in controls, the inclusion of pairwise interactions was still insufficient to predict the observed network patterns. The structure of activity patterns in control animals is therefore likely shaped by higher order interactions^22^ (triplet, quadruplet, etc.) that are either diminished or absent in APP/PS1 animals. Previous studies have identified the role that these higher-order interactions play in shaping population activity in sensory systems and implicated the local circuits that give rise to such higher-order interactions^13^; our results hint at their importance in dCA1 and the extent to which they may be especially vulnerable to Aβ pathology in APP/PS1 animals.

