## Supplemental Materials for "Altered dorsal CA1 neuronal population coding in the APP/PS1 mouse model of Alzheimer’s disease"

### Supplemental methods

#### Animals

All experiments were performed in accordance with the guidelines of the Institutional Animal Care and Use Committee (IACUC) at the University of Rochester. 4 APP/PS1 double transgenic mice and 4 control mice were used in this study. The APP/PS1 mice expressed chimeric mouse/human amyloid precursor protein (Mo/HuApp695swe)[1] and mutant human presenilin-1 (PS1-dE9)[2] under the control of the neuron-specific prion protein promotor element[3]. All mice were males aged 11 to 13 months. By this age, amyloid plaques are present in dCA1[4,5] and behavioral deficits manifest in multiple domains, including spatial memory, cognition, and anxiety[6–9]. This study was restricted to males, as it is known that there are sex differences in amyloid burden and cognitive changes in mouse models of Aβ pathology. For instance, female APP/PS1 mice show higher levels of amyloid plaques, as well as Aβ40 and Aβ42 peptides, than age-matched male mice[10]. Moreover, a recent study using the related Tg2576 mouse model, showed that the relationship between cognitive impairment and Aβ levels itself depends on sex, with female animals exhibiting poorer performance on a reference memory task than males with a similar amyloid burden[11]. Additionally, multiple aspects of hippocampal structure and function, including LTP induction[12], seizure threshold[13], dendritic morphology[14], and synapse density[15], fluctuate across the estrous cycle. To avoid the effects of these confounding factors, we restricted our study to males. However, given the evidence for different prevalence of AD among human males and females[16,17], studying sex-specific differences in neuronal population activity in the context of amyloid pathology remains an direction for future research beyond the scope of this study.

#### Surgery

Prior to surgery, mice were anesthetized with an inhaled 1-2% isoflurane mixture. The scalp was resected and a custom 3D printed head frame made from polylactic acid (PLA) was affixed to the dorsal surface of the skull using veterinary adhesive (Vetbond, The 3M Company,

Maplewood, MN, USA) and dental cement (Ortho-Jet Powder and Jet Liquid, Lang Dental Mfg. Co., Wheeling, IL, USA). A metal screw was also implanted in the skull to serve as an electrical ground. For 72 hours, mice received postoperative 0.03mL subcutaneous injections of 0.3mg/mL buprenorphine every 12 hours.

**Run-wheel training**

For 7 days following headframe implantation, mice were placed on a non-motorized running wheel in the recording rig for 1 hour daily. Mice were secured in place via clamps to the headframe, but they were free to run on the wheel. During these training sessions, no electrophysiological recordings were performed, although running behavior was recorded via a rotational encoder attached to the wheel. The purpose of these training sessions was to accustom the mice to running on a wheel in the head-fixed setup.

**Electrophysiology**

Following the training period, mice were anesthetized using 1-2% inhaled isoflurane and a craniotomy was performed over the right dorsal CA1 (dCA1) region using stereotactic coordinates (-2.5mm caudal 1.5mm lateral of bregma)[18]. Mice were then transferred to the running wheel and allowed to recover from anesthesia. A 4-shank, 128-channel nanofabricated silicon array[19] was vertically lowered into dCA1. While the mouse was awake and behaving on the running wheel, extracellular voltage recordings were collected at 30 kHz in the 500-3500Hz frequency band. Simultaneously, the instantaneous running velocity of the mouse was also collected.

**Spike identification and sorting**

All data analysis was performed in MATLAB (The Mathworks, Natick, MA, USA). To eliminate background activity, the mean signal from all 128 channels was subtracted from the signal from each individual channel. Intervals during which the voltage signal surpassed a threshold of 8 standard deviations (SD) above or below 0 were tagged as putative action potentials, or spikes. Putative spikes that were separated from other spikes by less than 1.3ms were eliminated, as they were difficult to sort. Subsequently, a raster of all putative spikes was produced and intervals containing high-voltage artifacts that manifested on nearly all channels were visually identified and eliminated. Some putative spike waveforms crossed the 8 SD threshold at both positive and negative voltages; these double-counted putative spikes were identified and consolidated. Putative spikes that remained at the end of these pruning steps were preserved as true spikes.

Next, spikes were assigned to different units, or putative neurons. Due to differences in intrinsic electrophysiological properties, distance from the channel, and orientation relative to channel, different neurons produce different characteristic spike waveforms across multiple channels. The waveforms of all spikes from each channel were concatenated with those from 8 neighboring channels and projected onto a low-dimensional space using principal components analysis (PCA). The dimensionality of this space was defined by the smallest number of principal components that, when combined, would explain > 80% of the variance in the waveforms. Spikes corresponding to different putative units appeared as distinct clusters. These were automatically identified using a mixture of Gaussians model[20].

The mean waveform of each putative unit across multiple channels was visualized (**Fig. 1b**,**e**) and manually evaluated by two reviewers. Putative units with noisy or symmetric waveforms, as well as those with fewer than 50 spikes, were discarded. Only the units that passed the evaluation of both reviewers were preserved for subsequent analysis. Additionally, groups of putative units with highly similar waveforms were merged. Finally, pairs of units from different channels that had highly overlapping spike times were tagged as likely originating from the same neuron; one of these was discarded. The end result of the spike sorting process was visualized with a raster plot, in which each row represents the spiking activity of a given unit (**Fig. 1c**,**f**). 235 units were identified in the control mice and 239 units were identified in the APP/PS1 mice. Details on the number of spikes and units eliminated at each step are shown in **Figure S1**.

Mean firing rates (FR) were decreased in the APP/PS1 group as compared to the controls (control: 1.65±3.32 Hz, APP/PS1: 0.80±1.49 Hz, $p < 10^{-4}$, one-sided Wilcoxon rank-sum test, **Fig. S15**), consistent with previous work showing the presence of hyperactive and hypoactive subpopulations of neurons in mouse models of AD[21–23].

**Running behavior**

The run-wheel was not motorized and its movement was controlled entirely by the mouse, which could run in either forward or reverse directions or remain stationary. Wheel movement was recorded by a 2-bit rotational encoder at 30kHz. This was converted to units of cm/s by averaging the velocity across a 150ms bin in time steps of 10ms (**Fig. 1c**,**f**). Epochs of running were denoted as intervals when the run velocity exceeded 1cm/s. All other times were denoted as stationary epochs.

**First order statistics**

The mean firing rate (FR) of each unit was calculated by dividing the total number of spikes generated by that unit by the duration of the recording. An inter-spike interval distribution was generated for each unit by calculating the time between all pairs of successive spikes from that unit. The mean and variance of the ISI were calculated from this distribution. Each of these quantities was also calculated separately for running and stationary epochs.

**Entropy**

The spike train of each unit was binarized over 10ms non-overlapping windows. If that unit spiked at least once in that interval, it was denoted with a *1*, and if it was silent throughout that interval, it was denoted with a *0* (**Fig. 2a**). To estimate the entropy for each animal, 1000 random subsamples of 10 units were generated (**Fig. S16**). In each subsample, the state of all 10 units at a given time was denoted as a pattern (these patterns are also called words[24–26]). The relative frequency of the $2^{10}$ patterns in each subsample was computed and used to generate a pattern probability distribution (**Fig. 2a**). $\sum_{i=1}^{1024} -p_i \, log_2 p_i$ was then used on the pattern probability distribution to calculate the entropy of each subsample. Entropy conditioned on running or stationary states was calculated by using spike trains taken only from running or stationary epochs, respectively (**Fig. 2c**).

**Maximum-entropy models**

Maximum-entropy models[24,27] were fit using the maxent_toolbox software[28]. First, the intrinsic spiking bias terms (**$h_i$**) and pairwise interaction terms (**$J_{ij}$**) were computed from the binned and binarized spike trains. The resulting model was then used to predict the probability of each pattern: $P(\sigma_1\sigma_2 \ldots \sigma_N) = \frac{1}{Z} e^{\sum_i h_i \sigma_i}$ for the independent firing model and $P(\sigma_1\sigma_2 \ldots \sigma_N) = \frac{1}{Z} e^{\sum_i h_i \sigma_i + \frac{1}{2}\sum_{i \neq j} \sigma_i \sigma_j}$ for the pairwise interactions model. $\sigma_i$ denotes the binary state of each neuron and Z denotes the partition function, which normalizes the pattern probability distribution. The model pattern probabilities were then compared with the empirical pattern probabilities (**Fig. 3a-d**) using the Kullback-Liebler divergence (KLD)[29]. The smaller the KLD, the better the agreement between the predicted and model probability distributions.

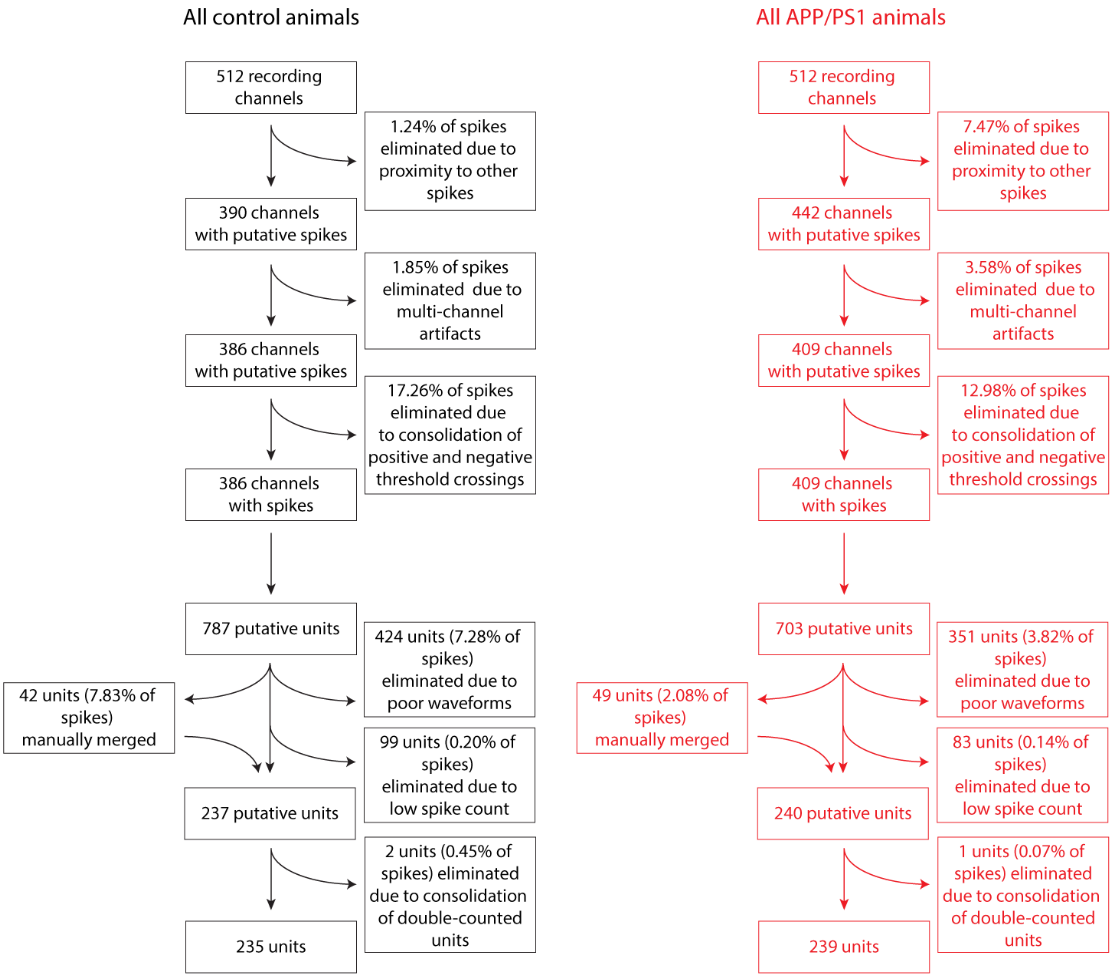


**Figure S1** Schematic of spike sorting pipeline, indicating the number and proportion of spikes eliminated at each step.

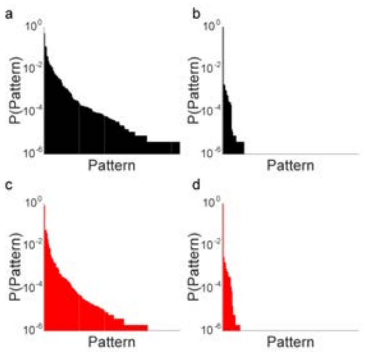


**Figure S2** Representative pattern probability distributions from a (**a**,**b**) control and (**c**,**d**) APP/PS1 animal. (**a**,**b**) Example of a high-entropy pattern probability distribution from a single 10-unit subsample in a (**a**) control and a (**c**) APP/PS1 animal. (**c**,**d**) Example of a low-entropy pattern probability distribution from a single 10-unit subsample in a (**c**) control and (**d**) APP/PS1 animal.

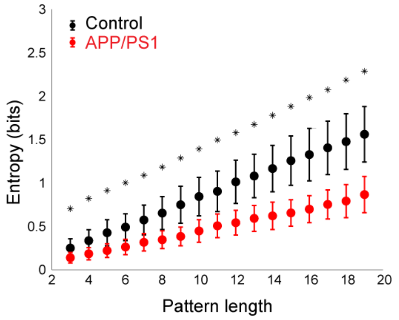


**Figure S3** Mean and standard errors of entropy estimates from each group as a function of the number of units used in the subsample. For both groups, entropy increases as pattern length increases, though the entropy of the APP/PS1 group is significantly lower than that of the control group for all pattern lengths tested ($p < 10^{-6}$, one-sided Wilcoxon rank-sum test, Bonferroni corrected).

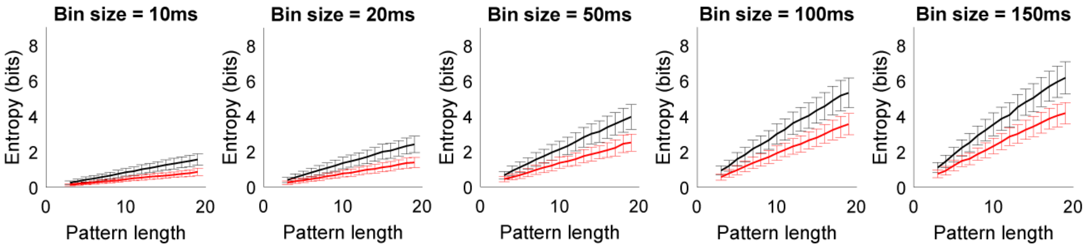


**Figure S4** The decreased entropy observed in APP/PS1 animals relative to controls is robust to choice of bin size and pattern length ($p < 0.05$, one-sided Wilcoxon rank-sum test, Bonferroni-corrected). Entropy in both groups increased with pattern length and with bin size. Error bars denote standard errors.

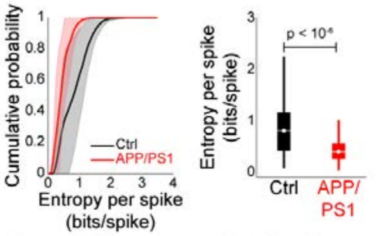


**Figure S5** Entropy normalized by firing rate (**a**) Cumulative histogram of entropy per spike from all animals in each group, with the mean denoted with the bold line and the standard deviations denoted by the shaded regions. A distribution of estimates is obtained by calculating the entropy of 1000 10-unit random subsamples. (**b**) Comparison of normalized entropy pooled across all the control (black) and APP/PS1 (red) animals. The APP/PS1 group had significantly lower entropy per spike than the control group ($p < 10^{-6}$, two-sided Wilcoxon rank-sum test).

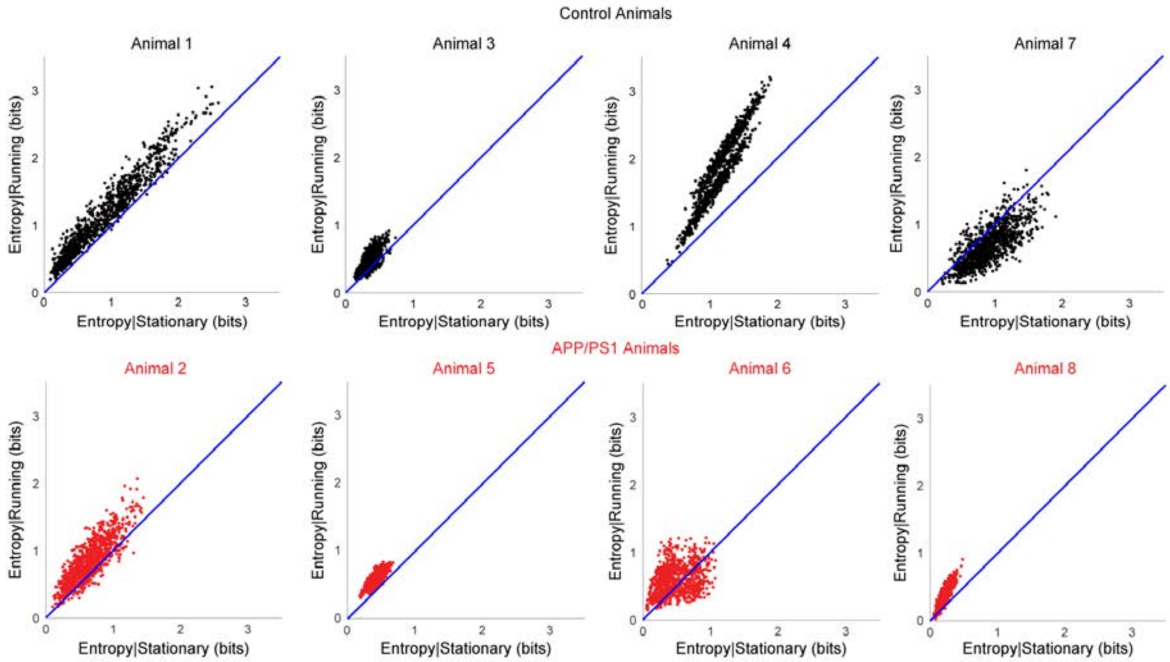


**Figure S6** Plots of entropy conditioned on running behavior for each animal. Each point denotes an estimate of conditional entropy based on a single 10-unit subsample of the neuronal population in that animal. Note that for most of the animals, both in the APP/PS1 and the control groups, the points are clustered above the unity line (blue), indicating that running is associated with increases in entropy.

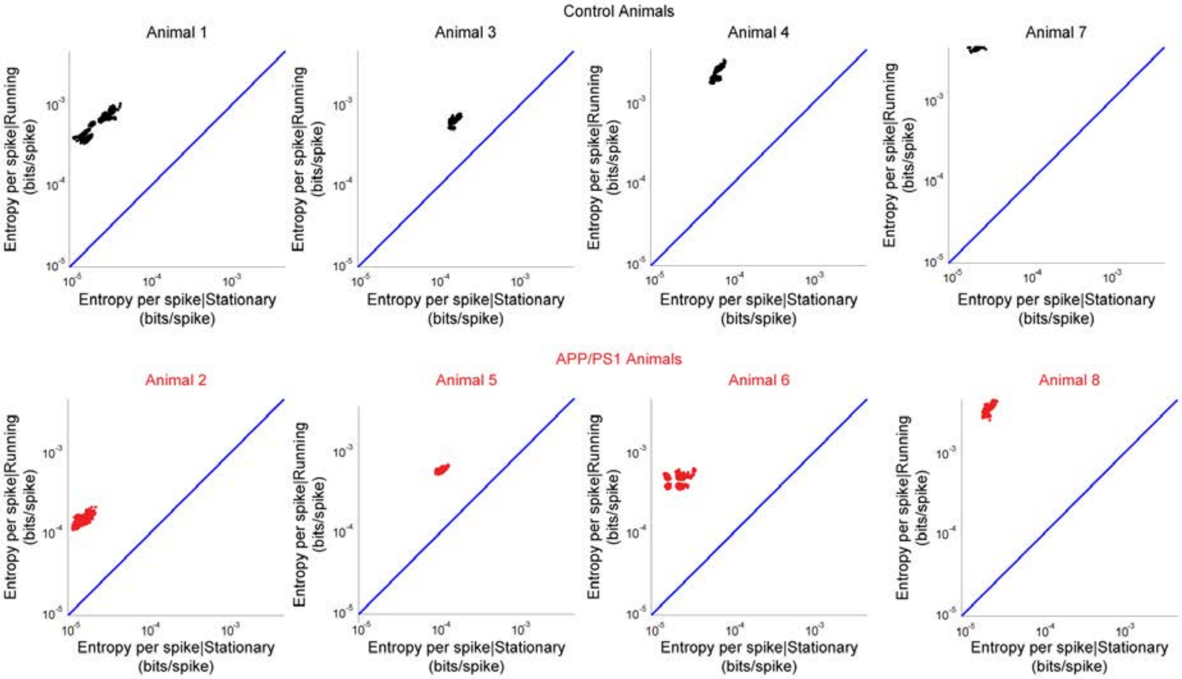


**Figure S7** Plots of entropy conditioned on running behavior for each animal, normalized by firing rate. Each point denotes an estimate of conditional entropy based on a single 10-unit subsample of the neuronal population in that animal. Note that for most of the animals, both in the APP/PS1 and the control groups, the points are clustered above the unity line (blue), indicating that running is associated with increases in entropy, even after normalizing by firing rate.

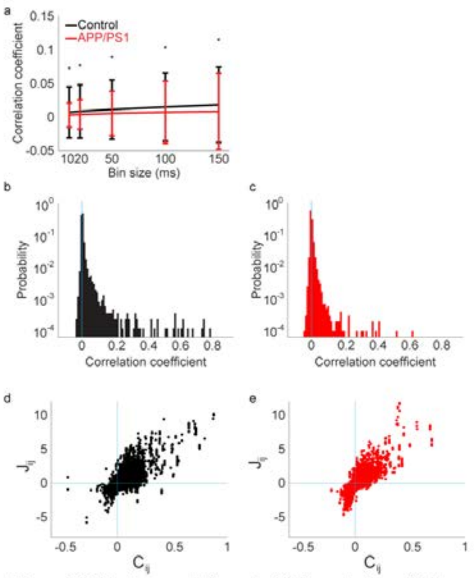


**Figure S8** Pairwise correlations in dCA1 populations. (**a**) Mean Pearson correlation coefficients of neuronal populations in control and APP/PS1 animals. For all bin sizes examined, correlations were larger for control animals than APP/PS1 animals (*$p < 0.05$*, Bonferroni corrected, one-sided Wilcoxon rank-sum test) (**b**,**c**) Histogram of Pearson correlation coefficients between neurons in all control (**b**) and APP/PS1 (**c**) mice. (**d**,**e**) Relationship between the Pearson correlation coefficient $C_{ij}$ and the maximum entropy model pairwise interaction term $J_{ij}$ for all pairs of neurons i and j for all control (**d**) and APP/PS1 (**e**) animals.

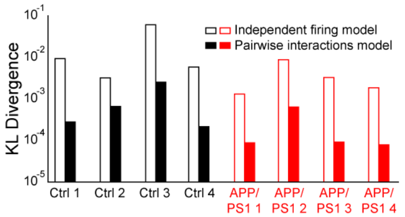


**Figure S9** Maximum entropy model fits for each animal. Empty bars show the KL divergence of the independent firing model and filled bars show the KL divergence of the pairwise interactions model. Control animals are in black and APP/PS1 animals are in red.

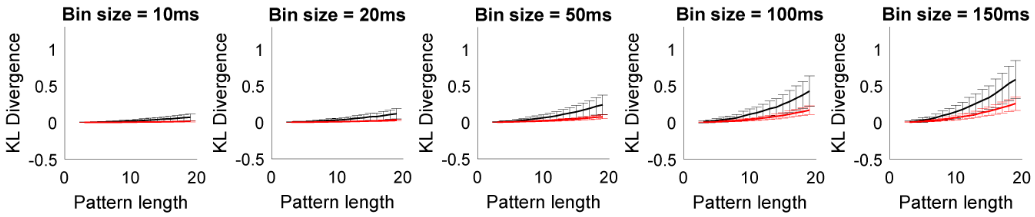


**Figure S10** The decreased KL divergence between the empirical and predicted pattern probabilities from the independent firing maximum entropy model observed in APP/PS1 animals relative to controls is robust to choice of bin size and pattern length ($p < 0.05$, one-sided Wilcoxon rank-sum test, Bonferroni-corrected). The KL divergence in both groups increased with pattern length and with bin size. Error bars denote standard errors.

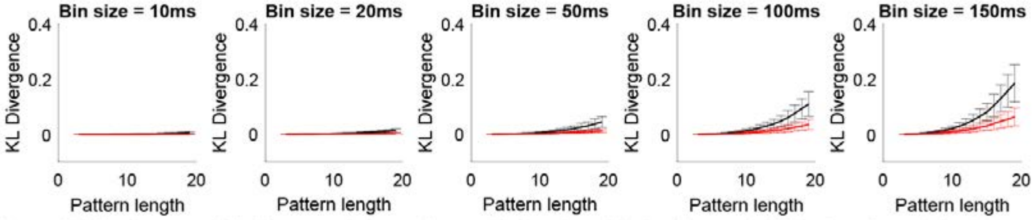


**Figure S11** The decreased KL divergence between the empirical and predicted pattern probabilities from the pairwise interactions maximum entropy model observed in APP/PS1 animals relative to controls is robust to choice of bin size and pattern length ($p < 0.05$, one-sided Wilcoxon rank-sum test, Bonferroni-corrected). The KL divergence in both groups increased with pattern length and with bin size. Error bars denote standard errors.

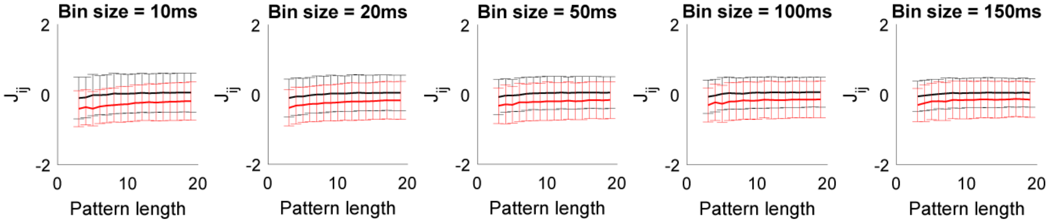


**Figure S12** The decreased maximum entropy $J_{ij}$ term observed in APP/PS1 animals relative to controls is robust to choice of bin size and pattern length ($p < 0.05$, one-sided Wilcoxon rank-sum test, Bonferroni-corrected). Error bars denote standard errors.

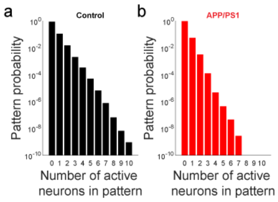


**Figure S13** Probability of patterns grouped by number of coactive neurons in (**a**) control and (**b**) APP/PS1 animals.

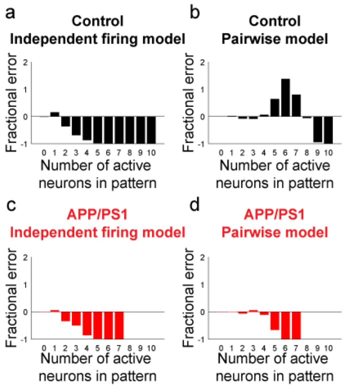


**Figure S14** Fractional error between predicted and empirical pattern probabilities, grouped by number of coactive neurons in each pattern for (**a**,**b**) control and (**c**,**d**) APP/PS1 animals. (**a**,**c**) Prediction error for the independent firing model. (**b**,**d**) Prediction error for the pairwise interactions model.

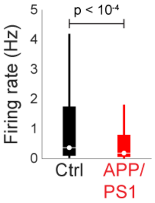


**Figure S15** Comparison of mean firing rates pooled across all control (black) and APP/PS1 (red) animals. The APP/PS1 group had significantly lower firing rates than control mice ($p < 10^4$, one-sided Wilcoxon rank-sum test).

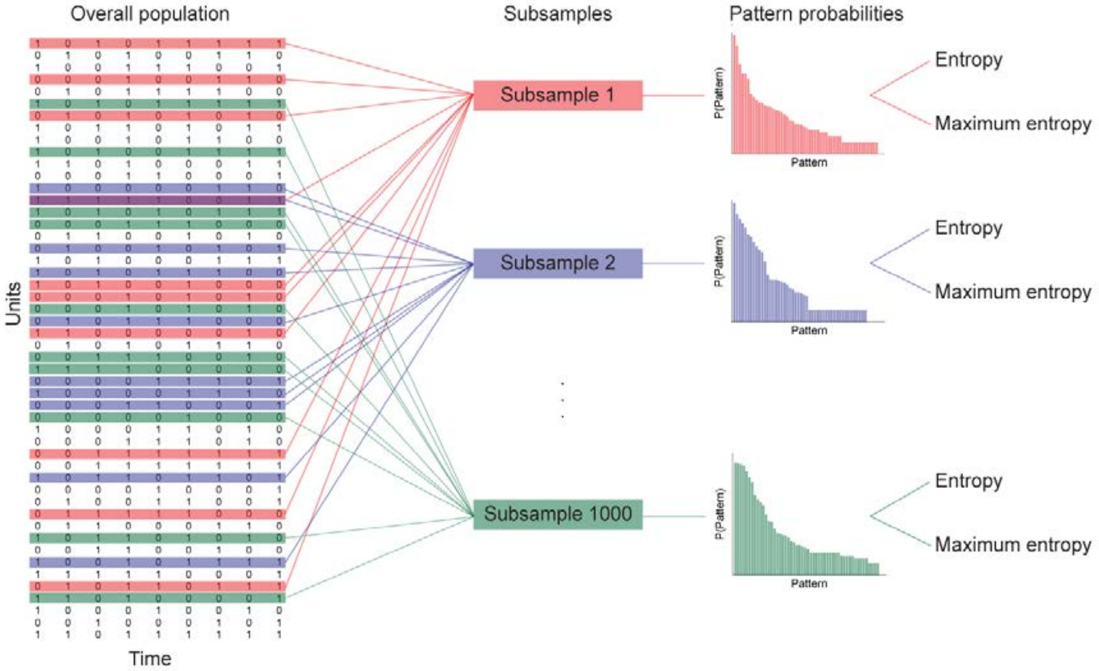


**Figure S16** Schematic of subsampling process. Subsets of neurons were repeatedly taken from the overall population to generate 1000 subsamples. For each subsample, pattern probability distributions were generated. In turn, the pattern probability distributions were used to estimate entropy and generate maximum entropy models.
